# Induced expression of *Xerophyta viscosa XvSap1* gene greatly impacts tolerance to drought stress in transgenic sweetpotato

**DOI:** 10.1101/603910

**Authors:** Wilton Mbinda, Christina Dixelius, Richard Oduor

## Abstract

**Key message Drought stress in sweetpotato could be overcome by introducing *XvSap1* gene from *Xerophyta viscosa*.**

Drought stress often leads to reduced yields and is perilous delimiter for expanded cultivation and increased productivity of sweetpotato. Cell wall stabilization proteins have been identified to play a pivotal role in mechanical stabilization during desiccation stress mitigation. They are involved in myriad cellular processes that modify the cell wall properties to tolerate the mechanical stress during dehydration in plants. This provides a possible approach to engineer crops for enhanced stable yields under adverse climatic conditions. In this study, we introduced the *XvSap1* gene isolated from *Xerophyta viscosa*, a resurrection plant into sweetpotato by Agrobacterium-mediated transformation. Detection of the transgene by PCR coupled with Southern blot revealed the integration of *XvSap1* in the three independent events. Sweetpotato plants expressing the *XvSap1* gene exhibited superior growth performance such as shoot length, number of leaves and yield than the wild type plants under drought stress. Quantitative real time-PCR results confirmed higher expression of the *XvSap1* gene in XSP1 transgenic plants imposed with drought stress. In addition, the transgenic plants had increased levels of chlorophyll, free proline and relative water content but malonaldehyde content was decreased under drought stress compared to wild type plants. Conjointly, our findings show that *XvSap1* can enhance drought resilience without causing deleterious phenotypic and yield changes, thus providing a promising candidate target for improving the drought tolerance of sweetpotato cultivars through genetic engineering. The transgenic drought tolerant sweetpotato line provides a valuable resource as drought tolerant crop on arid lands of the world.

## Introduction

Plants survival under adverse environmental conditions depends on integration of stress adaptive physiological and metabolic changes into their endogenous developmental systems. The world’s seventh important crop, sweetpotato [*Ipomoea batatas* (L.) Lam.] plays a significant role in food security and nutritional requirements, for millions of people in Asia and Africa (Bovell-Benjamin 2007). This crop also has an enormous potential to be commercially exploited as an industrial raw material (Kasran et al. 2015). Sweetpotato is widely cultivated on marginal lands due to its relatively high tolerance to abiotic stress. The orange-fleshed sweetpotato cultivars contain portentous β-carotene quantities which is a viable solution to combat vitamin A deficiency in sub-Saharan Africa (Laurie et al. 2018). The crop’s effective clonal propagation, its cultivation simplicity and high biomass generation make sweetpotato conceivably appropriate for molecular farming of valuable and novel products (Rukundo et al. 2017). Like other crops, sweetpotato is adversely affected by drought stress, which seriously hampers the crop productivity and negatively influence the expansion of sweetpotato cultivation (Jin et al. 2017). Additionally, as source of bio-energy, sweetpotato will mainly be grown on marginal land in the future. Improving the drought tolerance is essential for the crop’s adaption to the environmental stress and increasing productivity. Sweetpotato trait improvement through conventional breeding is constrained by its laborious and time-consuming genetics. These limitations principally arise owing to the crop’s high heterozygosity, high levels of male infertility, complicated polyploidy and production of few seeds because of its self-incompatibility, resulting to sturdy segregation of hybrid progenies and the loss of numerous valuable traits (Martin 1965, Mwanga et al. 2017). The fast advancement in plant biotechnology has unlocked new potentials for increasing tolerance to abiotic and biotic stresses to sweetpotato as well as improving its nutritional quality by identifying key genes and introducing them through genetic engineering Additionally, genetic engineering technology has the capability of introgressing genes from incompatible plant species or other organisms, an imperative phenomenon for crop improvement. In sweetpotato, little progress has been made in the generation and evaluation of transgenic events against drought tolerance.

Plant drought stress involve intricate regulatory processes that control water flux and cellular osmotic modification through the biosynthesis of osmoprotectants (Golldack et al. 2014). ‘Resurrection plants’ survive dehydration (losing over 90% of their water content) of their vegetative tissues to air dry state for extended periods and recover complete metabolic state after rehydration (Challabathula et al. 2016). These resurrection plants could serve as ideal models for ultimate design of important crops with enhanced stress tolerance. A representative of this special category of plants is *Xerophyta viscosa*. This species is a southern African native monocot species that has been extensively investigated to understand the genetic mechanisms of desiccation tolerance and may serve as potential plethora of superior genes that could introgressed into important crops (Costa et al. 2017). Of particular interest in this study is *XvSap1*, a stress-regulated protein, that was isolated from a cDNA library constructed from dehydrated *X. viscosa* leaves. A gene that has been associated with desiccation stress tolerance (Garwe et al. 2003). The XvSap1 protein has 49% identity to a cold acclimation protein WCOR413, WCOR413, from *Triticum aestivum* (Mohammadi et al. 2007). Earlier work on utilizing genes from *X. viscosa* have resulted in promising outcomes on drought stress improvement in important crops such as maize (Seth et al. 2016) and tobacco (Kumar et al. 2013) which encouraged us to exploit *XvSap1* for genetic engineering of sweetpotato.

The *XvSap1* gene is a highly hydrophobic protein. It encodes two membrane lipoprotein -lipid domains which are activated in leaves of *X. viscosa* during dehydration stress (Garwe et al. 2003). Although the function of *XvSap1* in *X. viscosa* is yet to be fully determined, there are three proposed hypotheses. First, *XvSap1* may be involved in stabilizing the plasma membrane; second, it may be involved in maintaining ion homeostasis; and last, *XvSap1* may be a G-protein-coupled receptors (GPCR) associated with signal transduction in osmotic stress (Iyer et al. 2008). G-protein-coupled receptors form a large family of proteins whose principle function is to transduce extracellular stimuli into intracellular signals (Kroeze et al. 2003). *Arabidopsis thaliana* and *Nicotiana tabacum* transgenic plants overexpressing *XvSap1* exhibited high tolerance to osmotic, salt, heat and dehydration stress (Garwe et al. 2006). In the present study, we generated and assessed transgenic sweetpotato plants expressing *XvSapI* under the control of its stress inducible endogenous promoter. The results demonstrate that transgenic sweetpotato plants expressing *XvSapI* had significantly improved tolerance to drought stress compared with the wild type plants.

## Materials and methods

### Vector construction for plant transformation

The full-length cDNA of *XvSap1* (accession no. CB330588.1), and its stress-inducible promoter was provided by Prof. Jennifer A. Thomson (University of Cape Town, South Africa), was cloned into the binary vector pNOV2819 at the *Bam*HI and *Hin*dIII restriction sites. The binary vector pNOV2819 contains the phosphomannose isomerase (*pmi)* gene as selectable marker that confers resistance to mannose. Since sweetpotato is not sensitive to mannose selection (Mbinda et al. 2015), the *pmi* was substituted with the neomycin phosphotransferase (*npt*II) gene flanked by the *nos* promoter and *nos* terminator at *Hin*dIII and *Kpn*I restriction sites. The binary vector harboring the transgene (*pSAP1-XvSap1*) and the selectable marker *npt*II, conferring kanamycin resistance, was mobilized into the disarmed *Agrobacterium tumefaciens* strain EHA105 via the freeze–thaw method (Weigel and Glazebrook 2006) and used for sweetpotato genetic transformation.

### Sweetpotato genetic transformation

Sweetpotato (*Ipomoea batatas* [L.] Lam., cv. Jewel (donated by International Potato Centre, Nairobi, Kenya) plants were used in this study. The plants were maintained *in vitro* by sub-culturing every four weeks at 27±1 °C under 16 h/8 h light/dark conditions (Mbinda et al. 2016). Explants were prepared from stem segments from 3 to 4-week-old *in vitro* plants by transversely cutting them into 6–10 mm segments before diving the segments along the axis. Genetic transformation, somatic embryogenesis and regeneration of putative transgenic plants were performed (Mbinda et al. 2018). Cefotaxime (250 mg/l) and kanamycin (50 mg/l) were supplemented to the medium to induce the development of transgenic cell clusters. The regenerated sweetpotato plantlets were transferred to soil and grown in a greenhouse at 27±1 °C.

### DNA extraction, PCR and southern blot analysis

Total genomic DNA was extracted from leaves of both wild type and putative transgenic sweetpotato lines using the CTAB method (Borges et al. 2009). The extracted genomic DNA was amplified using gene specific primers (Supp. Table S1) to evaluate to the presence or absence of *XvSap1*. Southern blotting was performed to evaluate the copy number and stable integration of transgene in the transgenic sweetpotato lines using standard techniques (Sambrook et al. 1989). Genomic DNA (20 µg) from PCR positive and wild type plant lines was digested with *Eco*RI. Agarose (0.8 %) gel electrophoresis was used to separate the digested DNA, before blotting onto a Hybond-N+ membrane, followed by UV cross-linking. A probe, amplified from the *XvSap1* plasmid (Supp. Table S2), was labeled with dCTP α-^32^P using Amersham Rediprime II DNA Labelling kit (GE Healthcare, Buckinghamshire, England) and used for hybridization at 42 °C. Blots were washed for 10 min at 42 °C and more stringently twice for 30 min at 55 °C. A third wash was done if the residual radioactivity on the membrane was considered high. Finally, the membrane was monitored using a Kodak Flexible Phosphor Imager (Eastman Kodak Co. New York, US).

### Drought tolerance analysis

Wild type and transgenic sweetpotato stem cutting (12 cm) were planted into pots containing mixed soil in greenhouse under 750-1000 µmol Em^−2^S^−1^ photosynthetic photon flux density with 28±2 °C, 16/8 day photoperiod and 80±5 % relative humidity for 4 weeks. Thereafter, and in a completely randomized design, two groups of plant treatments: normal conditions (61.7 % soil water content) and simulated water deficit stress (0.3 % soil water content) were set for 12 days. Morphometric parameters (shoot length and number of leaves), biochemical assays (chlorophyll, proline, malondialdehyde and relative water contents) and the *XvSap1* expression analysis of the different materials were evaluated. Three biological replications were prepared for each treatment. Further, transgenic and wild type sweetpotato plants were planted in pots and regularly irrigated for 30 days. Thereafter, the plants were exposed to drought stress for 90 days. Yield attributes in terms of fresh weight and dry matter were measured for comparative analysis. Control plants (wild type and transgenic plant lines) were irrigated throughout the experiment.

### Assessment of soil water content

Soil samples collected at 0, 3, 6, 9 and 12 days after withholding water were dried at 105 °C. The soil water content (SWC), as a measure of ‘drought’ severity was calculated using the weight fraction described by Coombs et al. (1987). SWC (%) = [(FW − DW)/DW] × 100; where FW was the fresh weight for portion soil from the internal area of sampled pots and DW was the dry weight for the soil portion after drying at 105 °C for 4 days.

### qRT-PCR analysis

Leaves from water stressed and control plants were collected at 0, 3, 6, 9 and 12 days after simulated drought stress. Total RNA was isolated using Spectrum Plant Total RNA Kit (Sigma-Aldrich, St. Louis, US). First strand cDNA was synthesized using gene specific primers (Supp. Table S3). Quantitative real-time PCR (qRT-PCR) was performed with Bio-Rad iQ5 l System Software 1.0. The sweetpotato ubiquitin gene was used as the internal control (Park et al. 2012). Relative expression values were analyzed with the comparative 2^−ΔΔCT^ formulae as described by Livak and Schmittgen (2001).

### Measurement of photosynthetic pigment

Chlorophyll contents, under normal and drought conditions, were measured in attached leaves of both wild type and transgenic plant lines with, SPAD chlorophyll meter (Minolta Co., Osaka, Japan), a non-destructive portable device. The youngest fully expanded leaf flag from the top of each plant were used. Measurement started 9 days prior to commencement of water deficit stress and continued for 12 days. The mean of the five values was taken per plant for each of three replications.

### Estimation of proline content

Colorimetric assay was used to analyse free proline content. Fresh leaf tissue (100 mg was homogenized in 10 ml of 3% (w/v) sulphosalicylic acid. The homogenate was filtered through number 1 Whatman filter paper. A 0.1 ml aliquot of the filtrate was mixed with 0.5 ml reaction solution [acidic ninhydrin [40% (w/v) acidic ninhydrin (8.8 µM ninhydrin, 10.5 M glacial acetic acid, 2.4 M orthophosphoric acid), 40% (v/v) glacial acetic acid and 20% (v/v) of 3%(v/v) sulphosalicylic acid]. The samples were incubated for 60 min at 100 °C and the reaction terminated by incubating the samples on ice for 5 minutes. Free proline from the samples was extracted by adding 1 ml of toluene and vortexing for 20 s. The absorbance at 520 nm was measured with UVmini-1240 UV-Vis Spectrophotometer (Shimadzu, Kyoto, Japan) using toluene as a reference. Proline content [µmol/g fresh weight (F. WT)] in leaf tissues was computed from a standard curve using 0-100 µg L-proline according to formula by Bates et al. (1973).

### Analysis of lipid peroxidation

Lipid peroxidation was measured in terms of malondialdehyde content, a product of lipid peroxidation, is an indicator of oxidative damage, caused by stress such as drought and salinity in plants (Del Rio et al. 2005). 100 mg of leaf tissue was homogenized by adding 0.5 ml of 0.1 % (w/v) trichloroacetic acid. The homogenate was centrifuged at full speed for 10 min at 4 °C. The collected supernatant was mixed with 0.5 ml of (1.5 ml 0.5% thiobarbituric acid diluted in 20 % trichloroacetic acid). The mixture was incubated 95 °C in water bath at for 25 min before ending the reaction by incubating on ice for 10 min. The absorbance was measured at 532 and 600 nm using spectrophotometer (UVmini-1240 UV-Vis) with 1% thiobarbituric acid in 20% trichloroacetic acid as control. The concentration of malondialdehyde (µmol/g FW) calculated as a measure of lipid peroxidation, was determined according to Health and Packer (1968).

### Determination of relative water content

Water loss in plants by transpiration was measured form a detached-leaf assay of the excised third leaf from wild type and transgenic sweetpotato plants. Fresh weight of the leaves was recorded immediately after excision and further incubated in a Petri plate at 28 °C in deionized water. The weight of these leaves was recorded after 1, 2, 3, 5 and 17 hrs. The rate of water loss was determined according to the formula described by Turner (1981).

### Statistical analysis

Analysis in all the experiments was carried out in three replicates. The data collected was analyzed using one-way analysis of variance and post hoc Fisher’s least significant difference test to determine significant differences between the means of each treatment. A value of at confidence level of 95% (p ≤ 0.05) was considered to be statistically significant. Minitab statistical computer software was used for data analysis. All graphs were drawn using SigmaPlot software.

## Results

### Sweetpotato transformation

A total of 167 sweetpotato stem explants were co-cultivated with *Agrobacterium* strain EHA101 carrying pNOV2819-XvSap1 binary vector (Supp. Fig. S1). The transformed stem segments gave rise to 133 kanamycin resistant calli. Three putative transgenic sweetpotato plants were regenerated from 117 surviving resistant calli (Fig 1A). The regenerated sweetpotato plantlets (Fig. 1B) were hardened for propagation in a biosafety greenhouse. Under normal conditions (25-27 °C and 61.7 % SWC), all transgenic sweetpotato lines were phenotypically and morphologically indistinguishable from the wild type lines (Fig. 1C and Fig. 1D).

**Fig. 1.**
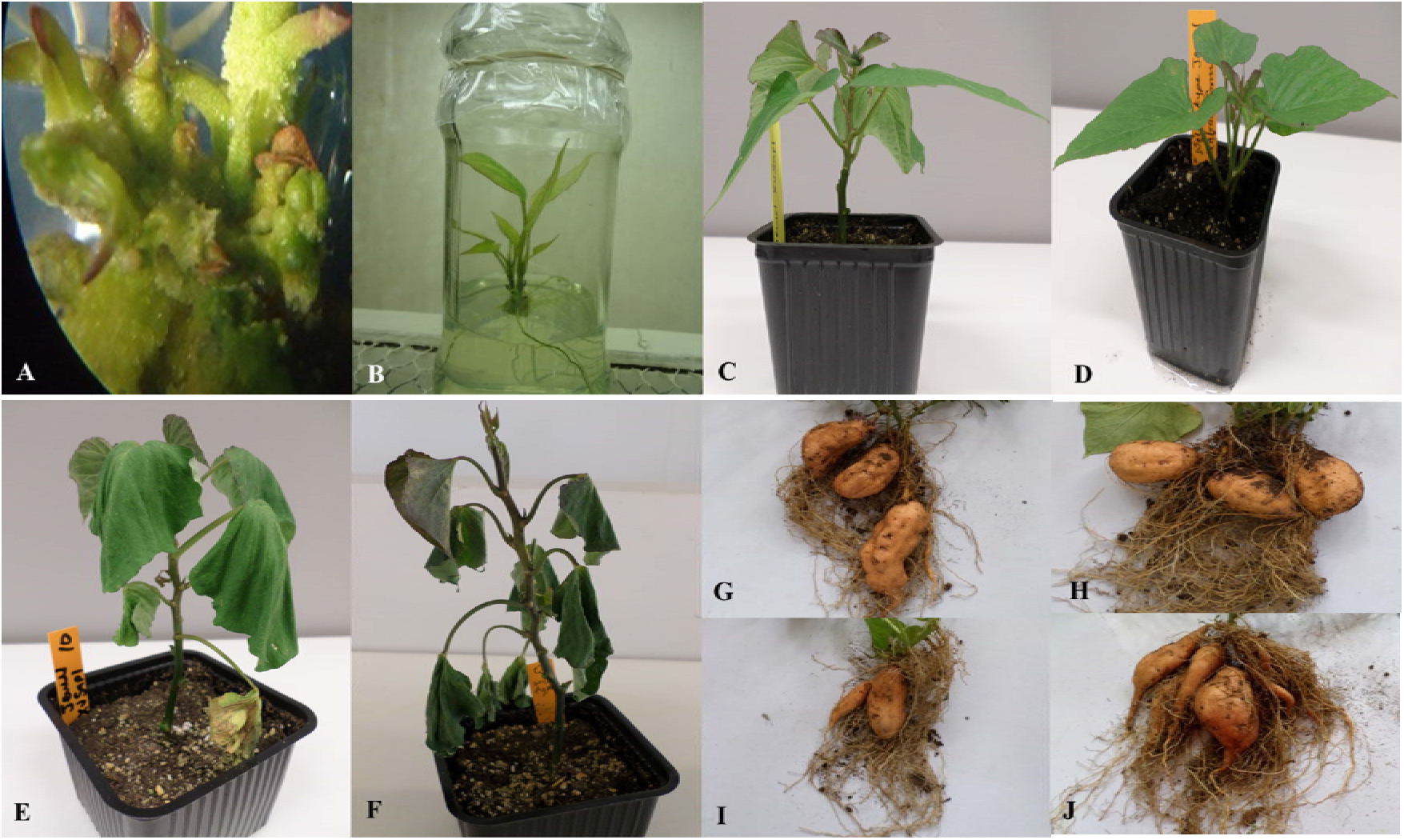
Transgenic sweetpotato lines obtained by somatic embryogenesis regeneration and responses of the drought stress in transgenic and wild type sweet potato plants. **A.** Developing embryo. **B**. Transgenic shoots obtained through somatic embryogenesis. **C** and **D**. morphological appearance of transgenic and wild type plants respectively. **E** and **F.** Phenotypes of transgenic and wild type plants grown in soil after 12 days of drought stress respectively. **G** and **H**. Yield performance of wild type and transgenic plants grown under normal conditions respectively. **I** and **J**. Yield performance of wild type and transgenic plants grown under drought conditions for 90 days respectively.

### Molecular characterization of transgenic sweetpotato plants

PCR analysis showed the presence of the 464 bp *XvSap1* fragment in the transformed plants but was absent in the non-transformed wild-type plants (Supp. Fig. 2). The results of Southern hybridization assay clearly showed all transgenic lines (XSP1-XSP3) contained a single copy insertion (Supp. Fig. 3). Further, results of reverse transcription-PCR (RT-PCR) analysis of the three transgenic independent events showed *XvSap1* mRNA transcripts, which was absent in the nontransformed control plants (Supp. Fig. 4).

### Phenotypic Characteristics

Under normal conditions, 4 weeks old transgenic and non-transformed controls sweetpotato plant lines did not have visible phenotypic differences (Fig. 1A and B). Our results therefore indicate that the introgression of *XvSap1* into sweetpotato did not lead to any major observable effects in plant architecture and growth properties. Upon exposure of the plants to severe water deficit stress (0.3 % SWC), shoot length of wild type control plants reduced by 55.2 % whereas that transgenic sweetpotato plant line was slightly higher (Fig. 2A). Similarly, after drought stress treatment, the average number of leaves in transgenic sweetpotato plants slightly declined by 15.7 % as compared to the wild type plants which had a drastic decline of 44.3% from an average of 8.7 leaves to 4.7 leaves (Fig. 2B). After 6 days of water stress (27.8 % soil water content), severe wilting was evidently observed in the leaves of the wild type plants (Fig. 1E) but of leaves of transgenic plants showed only minor damage (Fig. 1F). Severe dehydration symptoms were observed in the transgenic plants following 12 days of water deficit stress (0.3 % soil water content). The severe symptoms included wilted leaves, drying up of leaf edge and tip, and brown leaves. In both the wild type and transgenic plant lines, older leaves evidently displayed leaf wilting and chlorosis when the plants were exposed to severe drought stress (0.3 % soil water content). Overall, the wild type plants displayed much more severe symptoms compared to the transgenic plants.

**Fig. 2.**
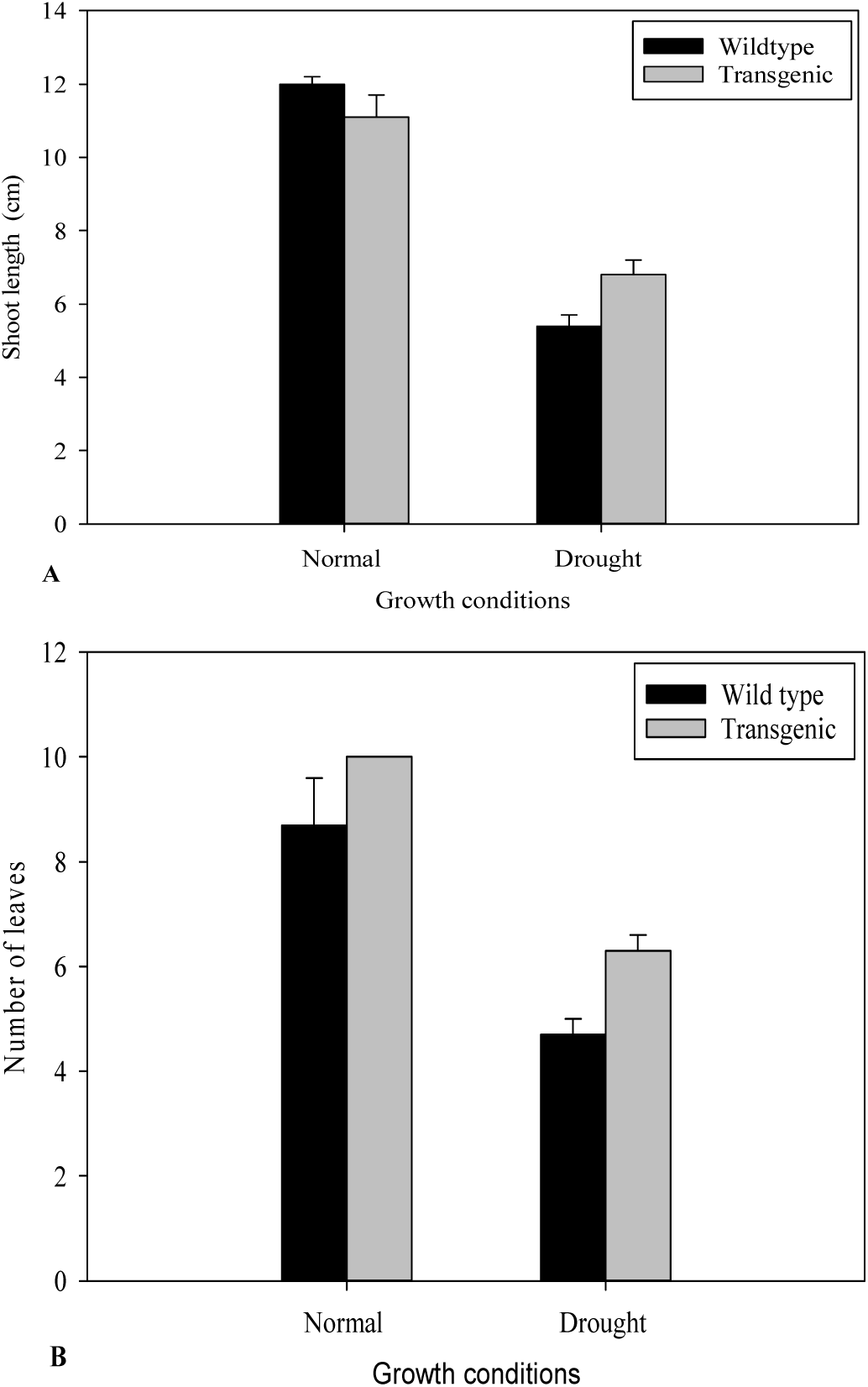
Phenotypic index changes of wild type and *XvSap1* transgenic sweetpotato plants under normal and drought stress condition. **A**. Effect of drought stress on shoot length. **B**. Effect of drought stress on number of leaves. Data are from three independent replicates of the same event.

Under normal conditions, there was no phenotypic difference in the tuber formation between the wild type (Fig. 1G) and transgenic plants (Fig. 1H). Following 90 days of drought stress, the yield of wild type plants (Fig. 1I) drastically reduced compared to transgenic plants that produced bigger, and higher numbers of tubers (Fig. 1J). When plants were regularly watered, the fresh weight yield and dry weight yield of both wild type and transgenic plants did not differ (Fig. 3A). Upon subjecting the plants to drought stress for 90 days, both the fresh weight and dry weight of transgenic and wild type plants were significantly changed. The fresh and dry weight yield per plant of transgenic plants under drought conditions was 341.5 g and 215.2 g respectively whereas wild type plants were 291.8 and 153.6 respectively (Fig. 2B). Further, we investigated the transcript expression levels in leaves of wild type and transgenic XSP-1 line under normal and drought stress conditions by qRT-PCR. The expression levels of XvSap1 in transgenic plants under normal condition was low but was induced under stress to 50-fold after 6 days of withholding water. Thereafter, expression started to decline, which is expected when using a stress-induced promoter (Fig. 4).

**Fig. 3.**
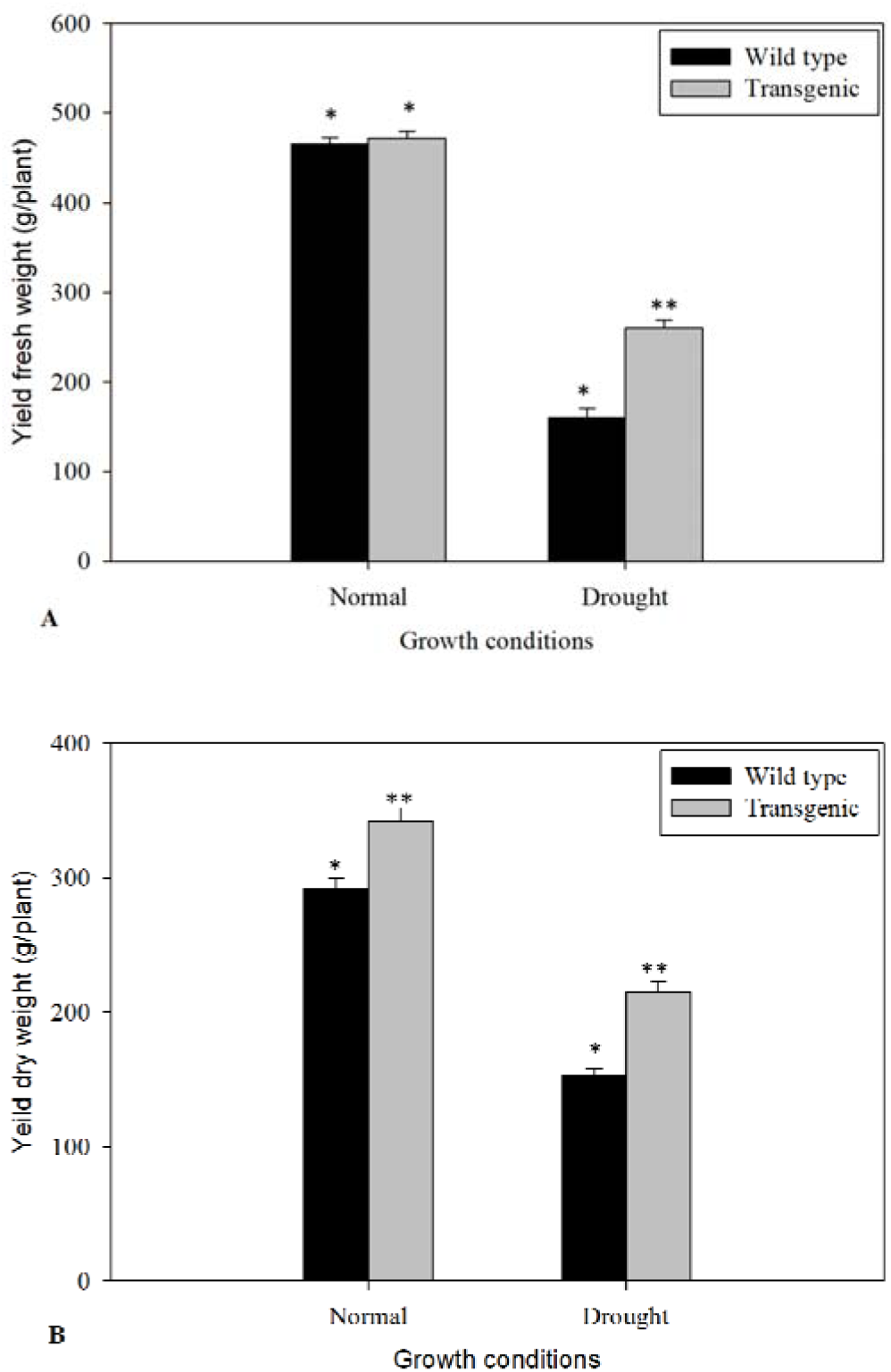
Yield attributes of tolerant transgenic plants and wild type plants grown in the drought stress. **A**. fresh weight **B**. Dry matter * and ** indicate a significant difference at p < 0.05.

**Fig. 4.**
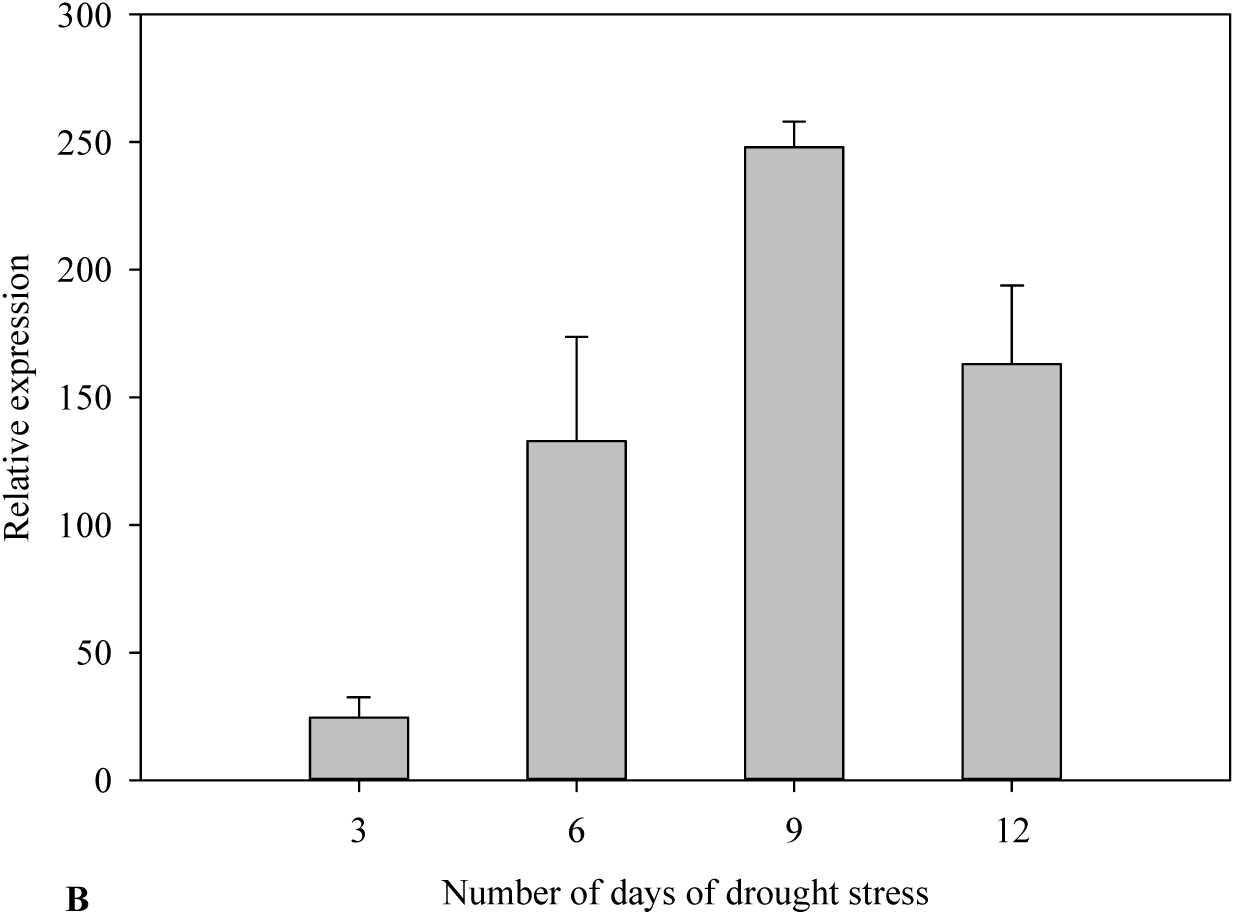
qRT-PCR analysis of *XvSap1* transcript levels in leaves of transgenic sweetpotato XSP1 line during 12-day water deficit stress.

### Biochemical characteristics

At the start of the experiment, chlorophyll content of both wild type and transgenic sweetpotato lines were the same and increased as the plants grew (Fig. 5A). Chlorophyll concentration in leaves of both wild type and transgenic plant lines progressively dropped with increasing drought stress intensity resulting in lower SPAD values compared to control treatment; plants grown under normal conditions throughout the experiment. In contrast, water deficit stress had mild effect on chlorophyll content in leaves of transgenic line compared to control, as shown in Fig. 5A.

**Fig. 5.**
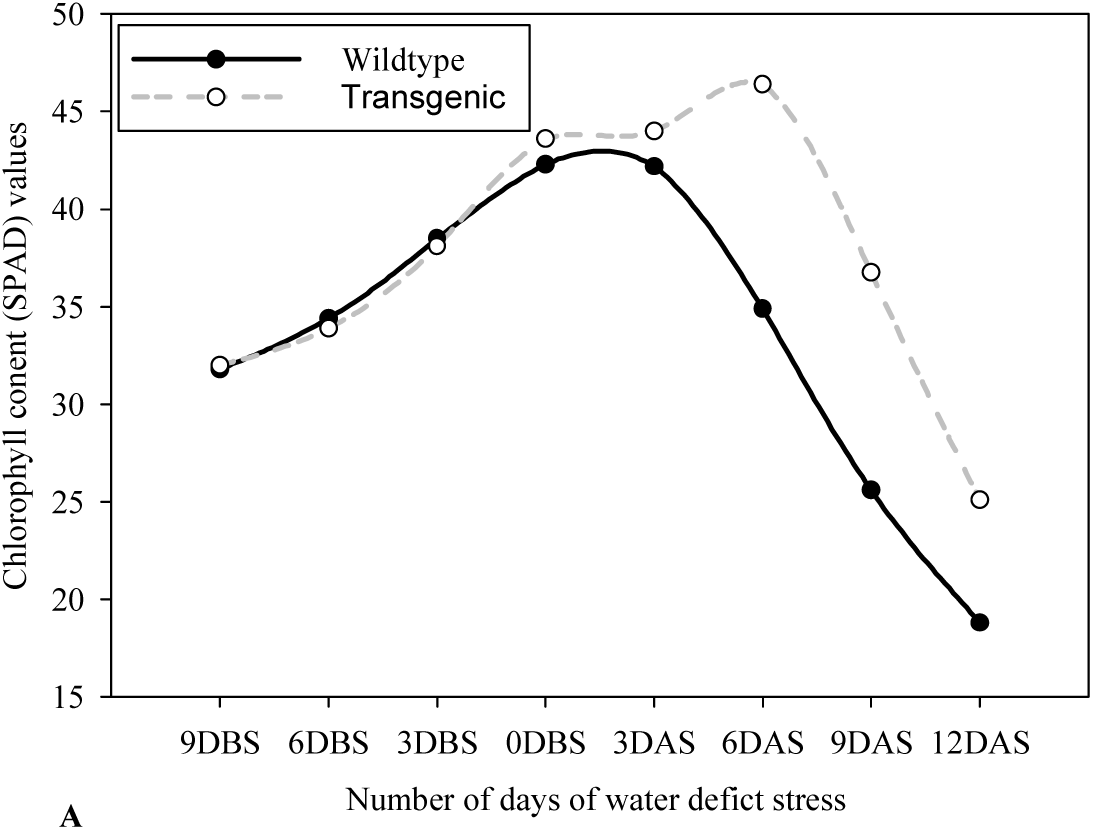
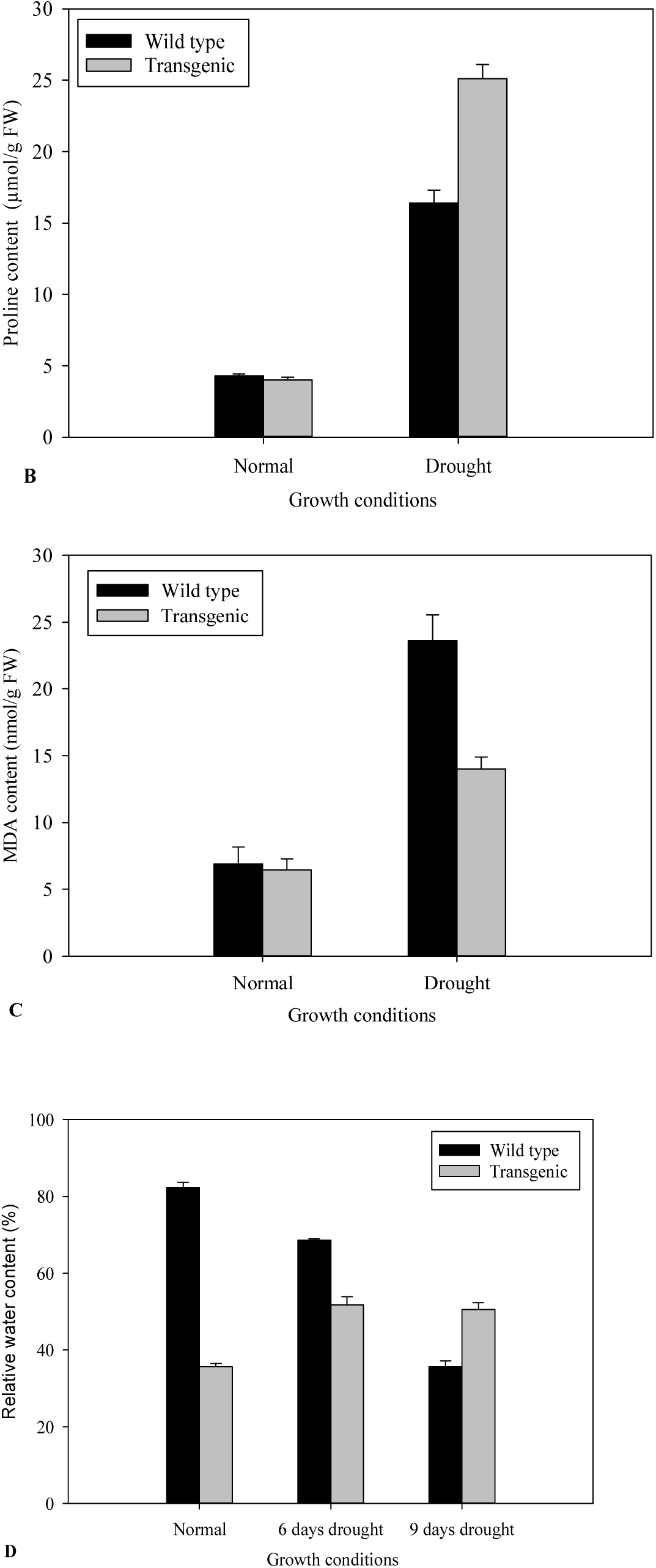
Biochemical index changes of wild type and XvSap1 transgenic sweetpotato plants under normal and drought stress condition. **A.** Effect of drought stress on chlorophyll content. Chlorophyll content was measured as SPAD values. **B.** Effect of drought stress on proline content. **C**. Effect of drought stress on malondialdehyde content. **D.** Effect of drought stress on relative water content. Data are from three independent replicates of the same event.

To evaluate the changes brought about by drought stress, we measured the accumulation of free proline in the shoot leaves of wildtype and the transgenic XSP1 line. Because level of free proline is known to be an indicator of drought stress resistance capacity in plants (Zandalinas et al., 2018). No significant difference in free proline concentration was observed in both wild type and transgenic leaves under normal conditions (Fig. 6B). After 12 days of water deficit stress experiments, the proline content in wild type plants increased 3.8-fold to 16.4 µg/g FW (Fig. 5B). In contrast, the XSP-1 line exhibited higher accumulation of proline x-fold??? compared to that of the wild type plants (Fig. 5B).

Further validation for the effect of *XvSap1* expression on lipid peroxidation was carried out by estimation of malondialdehyde (MDA) in the leaves. Under normal conditions, the levels found in the wild type line and the XSP1 line was similar (Fig. 5C). In contrast, imposition of water deficit stress to the plants resulted in about 3.4-fold increase in the malondialdehyde content in leaves of wild type plant line and much less malondialdehyde content in leaves of the transgenic sweetpotato plants was significantly lower than in the wild type plants (Fig. 5C).

### Relative Water Content

Under normal conditions, there were no significant differences in the leaf relative water content between control and transgenic sweetpotato plant XSP1 lines, and the relative water content of both lines was about 82 % (Fig. 6D). After subjecting the plants to drought stress for 6 days, the relative water content of the control wildtype leaves rapidly reduced by a 37.0 % to 51.8 %, while the relative water content of the transgenic line declined very gradually from 82. to 68.6 %, a 16.4 % decline (Fig. 5D). After 12 days of water deficit stress, the leaf relative water content of the transgenic plant had decreased by just 38.4 % as compared to 56.7 % in the wild type plants (Fig. 6D). These results demonstrate that the wildtype sweetpotato line was more drought sensitive than the transgenic line.

## Discussion

As drought frequency and severity are projected to increase in the future due to climate change perturbations, the ensuing deficit in water, especially in arid and semi-arid regions, will greatly reduce agronomic productivity. Increasing tolerance to these stresses through the development of crop varieties with improved tolerance is the only probable solution to extremely reduce the impacts of drought stress on important crops such as sweet potato crops. Through several studies, various stress tolerance genes that have profound effects in enhancing drought tolerance in plants have been identified and among them are the members of G-protein-coupled receptors (GPCRs) genes playing a crucial role in regulating plant water status under normal and adverse circumstances (Bai et al. 2018; Lu et al. 2018; Wong et al. 2018).

Numerous strategies have been applied to engineer plants with enhanced stress tolerance with limited success. One major problem is due to the undesirable or unfavorable phenotypic changes such as growth retardation and decrease in yield. Compared with wild type plants, we did not observe any phenotypic changes in transgenic sweetpotato plants. Similar results were also observed in our previous study when sweetpotato was genetically engineered with an aldose reductase, XvAld1, isolated from *X. viscosa* (Mbinda et al. 2018). Other studies using genes from *X. viscosa* and transferred into other crops; maize (Seth et al. 2016) and tobacco (Kumar et al. 20113) have also been successful, reaffirming the proposition that *X. viscosa* has prodigious potential for mining of valuable genes for biotechnological utilization with no nocuous variations to the accruing transgenic plants.

Under water deficit conditions, transgenic sweetpotato plants were identified as drought tolerant based on the reduction proportion of vine length and number of leaves compared to wild type. Upon re-watering after simulated drought treatment, transgenic plants resulted in rapid recovery with higher shoot height. Results from previous work corroborate our finding that dehydration stress in plant causes hydraulic restriction, with the ensuing high tension in the xylem water column and the stomatal closure of (Powell et al. 2017).

In plants, suitable amounts of reactive oxygen species (ROS) aggregation are vital for tolerance to abiotic stress (Das and Roychoudhury 2014). Nevertheless, excess ROS generation causes degradation of chlorophyll (Dietz et al. 2016). The degradation of photosynthetic pigments, which vital to plants principally for light harvesting during photosynthesis process and production of reducing power, is therefore one of the most sensitive indices and diagnostic tool for evaluation of plant responses to water deficit stress tolerance identification, genotypic variation, altitudinal variation of many crops such as potato (Rolando et al. 2015) and cassava (Orek et al.,2016). Plants can overcome drought stress by increasing the accumulation of chlorophyll which protects them by scavenging the excessive energy by thermal dissipation (Reddy et al. 2004). Waning of leaf chlorophyll concentration in response to drought stress is a common phenomenon in plants, occasioned by disordering chlorophyll synthesis and which results to changes in thylakoid membrane structure and plant chlorosis (Kalaji et al. 2016). When plants are subjected to environmental stresses, leaf chloroplasts are injured which leads to disrupted photosynthesis. Expression of *XvSap1* enhanced drought tolerance by stabilizing antioxidant status in leaf tissues during drought stress. We hypothesize that *XvSap1* enhanced drought tolerance by increasing stabilization of cell membranes of the desiccation stressed transgenic plants, The enhanced chlorophyll content exhibited by the transgenic line could be a result of stabilized antioxidant status in leaf tissues drought stress. Accumulation of proline and polyamines is a common response to various abiotic stresses and results in our study corroborate with responses of other crops to water deficit, such as kiwi fruit (Chartzoulakis et al. 1993), and cotton (Massacci et al. 2008), whose leaf chlorophyll content was higher in plants grown in normal conditions than under drought conditions.

Desiccation stress on plants induces osmotic and oxidative stress in plants, leading to cellular adaptive responses such as accumulation of compatible solutes (Chen and Murata 2011). Free proline is the most important osmolyte and signaling molecule which accumulates generally in the cytoplasm without prejudicing normal cellular physiological functions and also contributes in protection of membranes, proteins and enzymes against various stresses (Golldack et al. 2014). Proline accumulation has therefore a positive connection with their tolerance to various environmental stresses. The finger millet varieties with higher volume change when imperilled to drought stress have a higher proline content, but it is not clear if it is an adaptive response or a biochemical change as a result of the damage triggered by desiccation. Proline has been reported to contribute approximately 10-15% of the osmotic adjustments in water deficit stressed castor plants (Babita et al,.2010), pea cultivars (Sánchez et al. 1998) and transgenic sweetpotato (Mbinda et al. 2018) grown under drought stress. The biochemical relation between *XvSap1* gene action and the high proline content in the cells seems to be a sensitive balance

Enhanced reactive oxygen species (ROS) activity eventually leads to lipid peroxidation of the cell membranes and subsequent membrane leakage, a common phenomenon during various environmental stresses and causes severe damage to plant cells, leading programmed cell death. Malondialdehyde is a product of lipid peroxidation and its quantification is generally employed as an index of oxidative damages in plant tissues. The observed reduced lipid peroxidation in transgenic sweetpotato plants was probably due to lower reactive oxygen species generation under water deficit stress. These results suggest that inducible expression of *XvSap1* gene modulates oxidative stress responses. Moreover, plant growth and development are constrained by sstress factors such as scavenging and quenching of free radicals occasioned by ROS. These mechanisms are extensively regulated and plant tolerance to drought stress is related to their antioxidant scavenging ability. Enhanced levels of the antioxidant constituents impedes the stress damage before it develops to lethal status (Demidchik 2015; Mickelbart et al. 2015). Our results on growth and development of transgenic and control sweetpotato plant lines corroborate to what.

Ability to retain water during dehydration is an important strategy for plant tolerance to stress caused by drought. Consequently, measurement of relative water content is a rapid approach to estimate plant water status and provides projection levels of the cellular hydration levels after stress treatments (Sánchez-Rodríguez et al. 2010). Our study shows an enhanced water retention capacity in the transgenic sweetpotato plants. Decreasing transpiration rate is significant for plant survival under drought conditions.

Our results provided clear evidence that the presence of *X. viscosa* gene in transgenic sweetpotato plants displayed favorable traits associated with drought tolerance. From these results, we therefore believe that *XvSap1* gene may have great potential in sweetpotato crop improvements for drought stress.

## Supporting information

Supplememntary Matarial

## Acknowledgments

The authors thank Prof. Jennifer A. Thomson, University of Cape Town, South Africa for XvSap1 cDNA and XvPSap1 promoter and Prof Christina Dixelius for *npt*II. This work was funded by DST, India (Grant No. SR/SO/BB-37/2008). The authors acknowledge the Plant Transformation Laboratory, Department of Biochemistry and Biotechnology, Kenyatta University, Nairobi, Kenya, and the Department of Plant Biology, Swedish University of Agricultural Sciences, Uppsala, Sweden, for providing the laboratory facilities. This research work was funded by the National Council for Science and Technology, Kenya (Grant No. NCST/5/003/3rdSTICALL/109).

## Author contributions

RO, WM conceived and designed experiments. WM performed all experiments. RO and CD supervised the execution of the research, WM analyzed data and WM, CD and RO wrote the manuscript.

