## Supplementary material for "Induced expression of *Xerophyta viscosa XvSap1* gene greatly impacts tolerance to drought stress in transgenic sweetpotato": Supplememntary Matarial

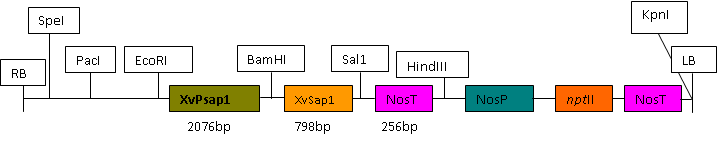

**Fig. S1** Plasmid vectors used in sweetpotato transformation. **RB,** right border of T-DNA; XvPSap1, stress inducible promoter from *X. viscosa*; *npt*II, neomycin phosphotransferase gene for plant kanamycin resistance; **NosT,** nopalin synthase terminator; XvSap1, truncated gene from *X. viscosa*; **LB,** left border of T-DNA.

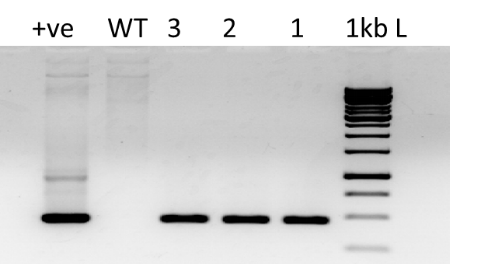

**Fig. S2**. PCR analysis of amplified transgene products from leaf tissues of the putative transgenic sweetpotato plants. Numbered lanes are transgenic plant lines, NT non-transformed plant for negative control, P vector for positive control and L 1-kb ladder (Thermo Fisher Scientific).

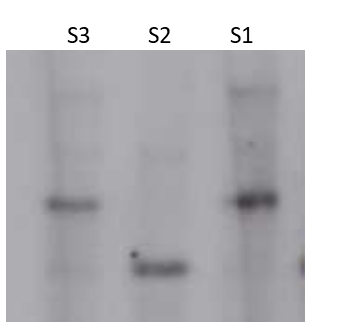

**Fig. S3.** Southern blot analysis of putative sweetpotato transgenics. 1. Southern blot showing integration of XvSap1 gene in transgenic plants. Digestion was performed using the *EcoRI* restriction enzyme. S1. 1, 2, and 3, are different XvSap1 transgenic sweetpotato lines.

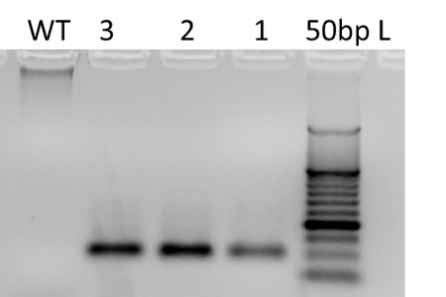

**Fig. S4** RT-PCR analysis confirming expression of *XvSap1* gene in leaves of transgenic sweetpotato plants. Numbered lanes are transgenic plant lines. *NT* non-transformed plant for negative control and *L* 50-bp ladder (New England Biolabs).

**Table S1 Primer sequences used for XvSap1gene amplification by PCR**

| **Primer** | **Sequence (5'- 3')** | **Product size (bp)** |
| --- | --- | --- |
| XvSap1-F | GCTAAACATGCGATCAAGCTCG | 464 |
| XvSap1-R1 | ATGATCCCGACGCTGTTTGAA |  |

**Table S2 Primer sequences used for synthesis of Southern blot probe**

| **Primer** | **Sequence (5'- 3')** | **Product size (bp)** |
| --- | --- | --- |
| cXvSap1-F | AATGAAGACCGACGTTGGAG | 413 |
| cXvSap1-R | CTCTGATTGTGTGGGCAATG |  |

**Table S3 Primer sequences used for cDNA amplification**

| **Gene** | **Primer** | **Sequence (5'- 3')** | **Product size (bp)** |
| --- | --- | --- | --- |
| XvSap1 | cXvSap1-F | AAGCTCTTCTTCCACCGACA | 103 |
|  | cXvSap1-R | CTCTGATTGTGTGGGCAATG |  |
| UBQ | cUBQ-F | CAAGCCGAAGAAGATCAAGC | 86 |
|  | cUBQ-R | GCACCTTTCCAGACTCATCC |  |
